# Exploring sex-specific alterations in early Alzheimer’s disease using network MRI analyses

**DOI:** 10.1101/2025.01.28.635128

**Authors:** Ines Ben Abdallah, Marion Sourty, Marion Rame, Laetitia Degiorgis, Amine Isik, Chantal Mathis, Laura-Adela Harsan

## Abstract

Alzheimer’s disease (AD) is characterized by the accumulation of amyloid-β plaques and tau neurofibrillary tangles, leading to progressive cognitive decline. Subtle cognitive and neural adaptations mark the preclinical stage of AD, occurring years before mild cognitive impairment and AD diagnosis. Throughout the continuum of AD, women (∼60% of AD cases) demonstrate faster rates of cognitive decline, greater hippocampal atrophy, and more extensive tau pathology compared to men. This is particularly evident in early AD phases. Therefore, investigating early sex-specific changes is crucial for identifying biomarkers and developing targeted interventions.

In the present study, we conducted a longitudinal investigation on the *App^NL-F^/MAPT* double knock-in (dKI) mouse model, to identify sex-specific resting-state functional connectivity (FC) patterns associated with early cognitive deficits. Male and female wild-type and dKI mice were tested for associative and long-term memory deficits, followed by resting-state functional magnetic resonance imaging to examine the default mode network (DMN) connectivity at 2 and 4 months of age. Female dKI mice exhibited earlier and more pronounced memory impairments, with deficits apparent at 2 months, while male deficits emerged at 4 months. FC analyses revealed distinct sex-specific alterations within DMN and between DMN hubs and memory processing nodes. Notably, female dKI mice showed hypersynchrony between the retrosplenial cortex (RSP) and key memory-related regions such as the entorhinal cortex (ENT) and hippocampus (HIP), but also towards subcortical regions overlapping thalamic nuclei, amygdala, and substantia nigra. Meanwhile, male dKI mice exhibited hypoconnectivity along RSP-HIP axis, RSP-reuniens nucleus, and RSP-sensorimotor cortex. Our data particularly highlight sexually dimorphic RSP-ENT and RSP-HIP connectivity.

These results underscore the critical role of sex in shaping neural network reorganization and cognitive decline during preclinical AD. These findings position neural network connectivity as a sensitive biomarker of early memory dysfunction and highlight the need for sex-specific investigations and potentially tailored therapeutic strategies in AD.

## Introduction

Alzheimer’s disease (AD) is a neurodegenerative disease characterized by brain accumulation of amyloid-β (Aβ) plaques and tau neurofibrillary tangles and a progressive loss of cognitive functions. The vast majority of AD patients develop the sporadic form around the age of 65 years. The onset and development of sporadic AD results from a complex interaction between non-modifiable factors such as age, sex, genetic risk factors like the apolipoprotein ε4 (APOEε4) allele coding for an apolipoprotein E variant, and modifiable factors such as lifestyle, environmental factors, and disease co-morbidities. Although increased longevity may explain why women represent over 60% of AD cases, they also show increased susceptibility to several risk factors such as AD comorbidities (e.g., depression and sleep disturbances) and deleterious effects of APOEε4 allele on cognition decline, especially after menopause (Rahman *et al*., 2019; Aggarwal and Mielke, 2023). In the early stages of AD, women show faster cognitive decline, reduced brain volume, and greater hippocampal atrophy and tau neuropathology (Lin *et al*., 2015; Koran *et al*., 2017; Buckley *et al*., 2018; Palta *et al*., 2021; Fernández *et al*., 2024). Progress in understanding the mechanisms underlying sex differences at the earliest stage of AD will be instrumental in the development of a ‘precision medicine’ approach in prevention, diagnosis, and treatment.

Less than 1% of AD patients develop between 30 and 60 years an autosomal dominant AD (ADAD) caused by genetic mutations in amyloid precursor protein (APP), presenilin-1 (PSEN1), or presenilin-2 (PSEN2) (Bateman *et al*., 2011). Because cognitive, neuropathological, and molecular features are similar to sporadic AD, investigations in ADAD mutation carriers have offered valuable insights in early biomarker changes in the presymptomatic stage of the disease (Jack *et al*., 2010; Bateman *et al*., 2012; van der Ende *et al*., 2023). This preclinical stage can precede for several years the prodromal mild cognitive impairment (MCI) stage of AD (Sperling *et al*., 2011). One of the major clinical symptoms in MCI is an episodic memory impairment affecting the recall of a specific event from one’s past in its unique spatial and temporal context (Tulving, 1983). This memory requires association and integration of the event’s « what », « where » and « when » dimensions, which rely heavily on the hippocampal formation and interconnected medial temporal lobe structures, such as the entorhinal cortex (ENT) (Sugar and Moser, 2019; Sakon and Kiani, 2022). The vulnerability of these regions to early AD pathology is coherent with the emergence of episodic memory defects (Van Hoesen and Hyman, 1990; Braak and Braak, 1991b, 1991a; Nyberg *et al*., 1996; Stoub *et al*., 2006). Cognitive evaluations of presymptomatic ADAD patients and older APOEε4 carriers showed deficits in short-term associative memory and accelerated forgetting in recognition tasks, indicating detectable impairments in specific cognitive functions during the preclinical stage well before overt symptoms appear (Parra *et al*., 2010; Sinha *et al*., 2018; Weston *et al*., 2018; Yang *et al*., 2021). Reduced performances in these tasks have also been linked to preclinical signs of AD in cognitively normal elderly (Rentz *et al*., 2011; Stark *et al*., 2013; Yeung *et al*., 2019). Early alterations in brain networks supporting associative and long-term memories may, therefore, contribute to episodic memory fading in the course of AD (Grande *et al*., 2021).

The progression of AD from the preclinical phase to dementia emphasizes the importance of detecting early neural changes that drive subtle cognitive and behavioral alterations emerging before the onset of overt clinical symptoms. Understanding these early changes is critical for identifying the neural mechanisms underlying AD’s initial stages and for developing targeted interventions. In this context, exploring alterations in brain networks that support associative and long-term memory is particularly relevant, as these networks are among the first to exhibit dysfunction in biomarker-positive individuals and those at genetic risk for AD.

To investigate these early network changes, advanced neuroimaging techniques like resting-state functional magnetic resonance imaging (rs-fMRI) (Biswal *et al*., 1995, 2010) offer a powerful, non-invasive approach to exploring network organization in the early phases of AD. By capturing spontaneous brain activity when no specific task is performed (Fox and Raichle, 2007; Raichle, 2011; Betzel *et al*., 2014; Chuang and Nasrallah, 2017), rs-fMRI enables the identification of functional connectivity (FC) patterns across the whole brain. These networks reflect the brain’s functional organization and consistently correspond to well-known cognitive domains (Greicius *et al*., 2003; Raichle, 2011, 2015b). Among them, the default mode network (DMN) is a key resting-state brain network (Raichle, 2015a), shown to be involved in various cognitive functions involving memory (episodic, spatial, semantic), self-reference, and social cognition (Menon, 2023). It is most active during rest and shows reduced activity during task performance or goal-oriented behavior. Including regions such as the dorsal and ventral prefrontal cortex, posterior cingulate cortex, precuneus, and inferior parietal cortex (Raichle *et al*., 2001; Raichle, 2015a), the DMN plays a critical role in cognitive processes, including episodic memory processing (Buckner *et al*., 2005). Alterations in DMN have been closely associated with neurological disorders like AD and cognitive decline (Greicius *et al*., 2004; Dillen *et al*., 2016; Wang *et al*., 2018). Emerging research also highlights the influence of factors like age and gender in the modulation of DMN connectivity, revealing changes across the lifespan and potential sex-specific features of this network organization (Wang, Therriault, Servaes, Tissot, Rahmouni, Macedo, Fernandez-Arias, Sulantha S. Mathotaarachchi, *et al*., 2024).

Independent component analysis (ICA) and seed-based analyses are two complementary methods for studying brain FC using rs-fMRI data. ICA uncovers independent brain networks (Calhoun *et al*., 2001) and has been instrumental in identifying differences in the DMN between men and women with AD (Damoiseaux *et al*., 2012; Caldwell *et al*., 2019; Pereira *et al*., 2020). Moreover, brain regions shared by the DMN and memory circuitry were found to be highly sexually dimorphic (Spets and Slotnick, 2022; Spets *et al*., 2024). Early midlife women exhibit for instance greater DMN connectivity between the left and right hippocampus (HIP) than men (Spets, Fritch and Slotnick, 2021; Spets *et al*., 2024). Moreover, within the female population, only postmenopausal women showed a loss of the ability to reduce left– right hippocampal connectivity within the DMN during memory encoding, a pattern linked to poorer memory performance (Spets *et al*., 2024). Additionally, seed-based analysis, which targets specific brain regions, has revealed altered hippocampal connectivity to the precuneus and brainstem in women with MCI (Williamson *et al*., 2022, 2024). In an amnesic subtype of MCI, decreased connectivity in the right HIP within the DMN was observed (Wang *et al*., 2019).

Functional brain imaging using similar methods in animal models has identified comparable networks (Zerbi *et al*., 2015; Grandjean *et al*., 2020; Mandino *et al*., 2021; Zerbi, 2022). DMN-like patterns are currently considered conserved across species (Zerbi, 2022; Pagani *et al*., 2023). These studies reinforce the value of such models in understanding brain connectivity and provide key insights into the neural underpinnings of health and disease (Zerbi *et al*., 2014; Grandjean *et al*., 2016; Degiorgis *et al*., 2020; Mandino *et al*., 2021, 2024). However, few studies have integrated fMRI approaches in animal models to comprehensively examine how sex influences functional network patterns in preclinical AD in relationship to behavioral assessments. Limited animal research in this area highlights the need for a more thorough understanding of how male and female brains may exhibit distinct patterns of network organization and their disruption in the early development of an Alzheimer’s pathology.

To address this gap, the present longitudinal study of the *App^NL-F^/MAPT* double knock-in (dKI) mouse model (Saito *et al*., 2019) was designed to identify sex-specific resting state network connectivity signatures associated with the onset of cognitive deficits similar to those found in preclinical AD. A cohort of male and female wild-type (WT) and dKI mice was evaluated in associative memory and long-term recognition tests before rs-fMRI was used to investigate their DMN at the ages of 2 and 4 months. These ages were selected as the dKI model develops key features of AD from early amyloid and tau markers, and associative memory deficits reminiscent of the preclinical stage at 4 months (Borcuk *et al*., 2022) before gradual increase in neuropathology leading to Aβ deposition, neuroinflammation, and increased tau phosphorylation at 24 months (Saito et al., 2019).

In this work, female dKI mice exhibit earlier onset of associative memory and long-term object recognition deficits, which emerge as early as 2 months of age in females, and 4 months in males. FC analyses further revealed significant sex-specific patterns, particularly within the DMN and its interaction with memory-related regions, in dKI mice and their WT controls. These differences highlight the critical role of sex in shaping the trajectory of cognitive decline in the preclinical phase of AD, pointing towards neural network reorganization as a sensitive biomarker of early memory dysfunctions.

## Materials and Methods

### Animals

The *App*^NL-F^/*MAPT* dKI and WT mouse line were initially derived at the LNCA from a 3-step breeding strategy initiated with heterozygous single knock-in *App*^NL-F^ mice and microtubule-associated protein tau (*MAPT)* mice obtained from the RIKEN BioResource Center in Japan (Saito *et al*., 2014; Hashimoto *et al*., 2019). Every fourth generation the mouse lines were backcrossed on their C57BL/6J background. The *App^NL-F^* gene harbors K670N/M671L (Swedish) and I716F (Iberian/Beyreuther) ADAD mutations within a humanized Aβ sequence replacing the murine fragment. The human *MAPT* gene replaces the murine fragment and enables the expression of the six tau isoforms. Magnetic resonance imaging (MRI) experiments enrolled four groups of mice (n=17/group): dKI female and male groups and WT female and males. Mice were group-housed (2 or 3 per cage) at the age of 4-5 weeks in a 12/12-hour light/dark cycle, with food and water ad libitum. They were evaluated in memory tasks and then imaged 2 weeks later at the age of 2 and 4 months (see Supplemental Fig. S1, experimental design). One WT male died before the 4-month scanning. Experimental protocols were approved by the regional committee of ethics in animal experiments of Strasbourg (CREMEAS: APAFIS#25628, APAFIS#27470).

### Memory tasks

Spontaneous object exploration tasks were used to detect early impairments in recognition memory. The mice were tested at 2 and 4 months for long-term object recognition in the Novel Object Recognition (NOR) task with a 24-hour delay interval and short-term associative memory in the Object-in-Place (OiP) task with a 5-minute delay. Experiments took place during the light phase in a 92.5*92.5*35 cm open field. Habituation and experimental procedure are detailed in Borcuk et al. (2022). Object exploration was recorded when the animal points its nose to the object within 2 cm.

### Long-term NOR task

Mice explored two identical objects during a 10-minute sampling trial the first day and were returned to their home cage for 24 hours. On the second day, one of the familiar objects was replaced by a new one, and the mice were given a 6-minute test trial. Performance was evaluated using a memory index: time on novel object - time on familiar object / total time exploring both objects (stated N-F/N+F on graphs).

### Associative OiP memory task

Mice explored two different objects during a 10-min sampling trial and were maintained in a holding cage for 5min. The object was then replaced by two copies of one of these objects, and the mice were given a 6-minute test trial. Memory index: time on new object-place association – time on familiar object-place association / total time exploring both objects (stated N-F/N+F on graphs).

### Rs-fMRI experiments

#### Animal preparation for MRI

During MRI acquisition, all animals were monitored for respiration rate and body temperature. A heating system via water pipelines implemented into the mouse bed maintained the body temperature at 37 ± 1°C during scanning. Anesthesia was induced with isoflurane 3% for 4 minutes and then reduced to 0.5% when injected with a medetomidine bolus subcutaneously (0.1mg.kg^-1^). During rs-fMRI acquisitions, mice were anesthetized by a combination of isoflurane (0.5%) and medetomidine (0.1mg.kg^-1^ bolus sc). Anatomical T2-weighted data was further collected under 1.5% isoflurane. After the brain MRI session, the mice were injected with 0.1mg.kg^-1^ glucose NaCl and allowed to spontaneously recover.

#### MRI data acquisition

Brain imaging was performed using a 7T small bore animal scanner (Bruker BioSpec 70/30 Germany) and a ^1^H four-channel phased array receive-only MRI mouse brain CryoProbe combined with a volume transmission coil 86mm diameter and the ParaVision software version 6.01 (Bruker, Germany). Rs-fMRI data was acquired with a GE-EPI sequence – TE/TR=15ms/2137ms; resolution=0.14×0.18×0.35mm^3^; 43 interleaved slices of 0.35 mm thickness and 500 volumes/ in 17min acquisition time. Anatomical images were acquired with a Turbo RARE T2-weighted sequence – echo time (TE)/repetition time (TR)=20ms/6000ms; resolution=0.07×0.07×0.35mm^3^/11min.

#### MRI data processing

Preprocessing of rs-fMRI data was performed using SPM12, ANTs, and in-house scripts in Matlab. Data underwent slice timing correction (SPM12), intensity bias correction (N4BiasFieldCorrection ANTs) (Avants, Tustison and Song, 2009), interslice drift correction, motion correction, spike sorting (Mazaika *et al*., 2009), and spatial normalization to group template in the Allen Mouse Brain Atlas space. A 0.45mm FWHM smoothing (SPM12) and band-pass filtering were applied to select blood oxygen level-dependent signal frequencies between 0.01 Hz – 0.22 Hz. For all rs-fMRI analyses except ICA, a least square approach was used for the regression of cerebral spinal fluid signal and 24 motion (Friston *et al*., 1996).

### Data analysis

#### Behavioural data

Memory indexes were analyzed with Prism 10 software. We used a two-way ANOVA with genotype and sex factors, followed by Fisher’s LSD test. A one-sample t-test was used to test within-group performance against chance level (memory index = 0).

#### Independent Component Analysis

A group-level spatial ICA was performed using the GIFT toolbox (Calhoun *et al*., 2001) to extract common resting-state networks (RSNs) from all mice. We used the implemented 2-step procedure with 170 components for the PCA and then we tested 15, 30, 50, and 100 for the ICA. We present here DMN-like rs-fMRI patterns from 30 ICA and 50 ICA. The Infomax ICA algorithm (Bell and Sejnowski, 1995) was repeated 10 times with ICASSO (Himberg, Hyvärinen and Esposito, 2004) for stable independent components. Each non-artefactual RSN spatial map and its associated time course were back reconstructed at individual level (Calhoun *et al*., 2001). Inter-group statistical analysis was conducted on back-reconstructed component maps to evaluate changes in connectivity patterns. Statistical maps are thresholded at T>2.6, p<0.01 uncorr, with cluster size ≥ 20 voxels.

Additionally, incidence maps were generated to display the spatial distribution of a component, and within each group, the percentage of animals displaying the pattern in each voxel (Supplemental Fig. S2).

#### Seed-based voxel-wise analysis

FC analysis was performed using the retrosplenial cortex (RSP) - key node of the DMN - as the seed region. The masks for the seed were extracted from Allen Mouse Brain Atlas. Spearman’s correlation was performed between the mean time course of RSP and all the other voxels in the brain to identify individual and group-specific RSP functional network. The inter-group differences in connectivity of RSP were assessed using a full factorial design in SPM to contrast the following groups: Females WT vs. Males WT (at 2 and 4 months); Females dKI vs. Females WT (at 2 and 4 months); Males dKI vs. Males WT (at 2 and 4 months); Females dKI vs. Males dKI (at 2 and 4 months). Within this full factorial design, we also included the Sex-Genotype interactions at 2 months and 4 months. The statistical thresholds were set at p < 0.02 - uncorr., with a minimum cluster size of 20 voxels.

## Results

### Early memory impairments emerge in a sex-dependent manner in dKI mice

To evaluate memory functions in dKI mice, we first assessed short-term associative memory performance in the OiP task with a delay of 5 min between sample and test trials (Fig. 1A), and then long-term object recognition memory performance in the NOR task with a 24h hour delay between sample and test trials (Fig. 1B).

**Figure 1:**
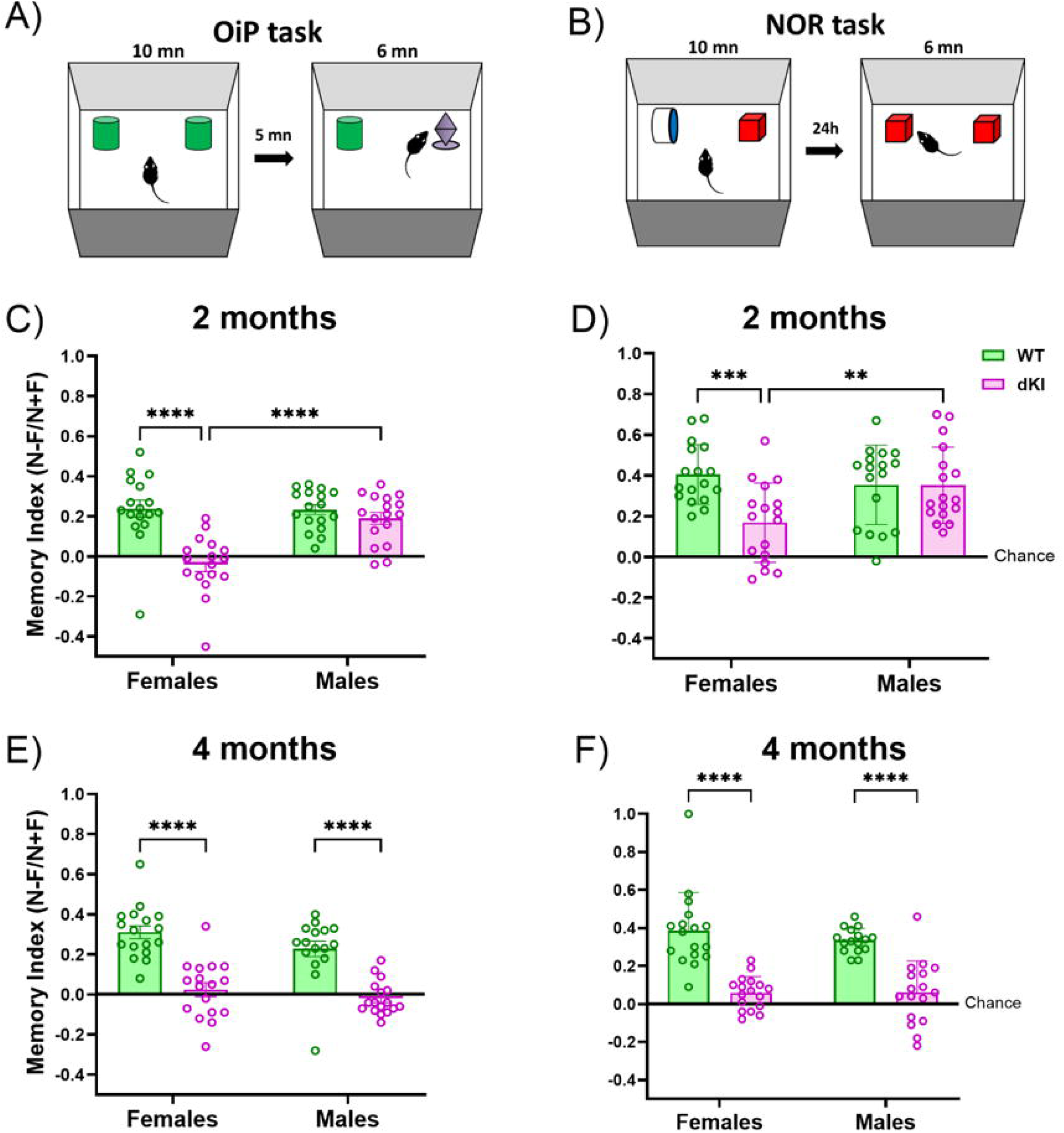
OiP and NOR performances of WT and dKI females and males at the ages of 2 and 4 months. (A) Schema of the OiP test protocol. OiP memory index for WT and dKI females and males at the ages of 2 months (B) and 4 months (C). (D) Schema of the NOR test protocol. NOR memory index for WT and dKI females and males at the ages of 2 months (E) and 4 months (F). WT females (n=17), dKI females (n=17), WT males (n=16), dKI males (n=17). Memory index performances are presented as mean ± SEM. Data were analyzed using a two-way ANOVA, followed by Fisher’s LSD test: **** p < 0.0001, *** p < 0.001, ** p < 0.01 for group difference.

At the age of 2 months, memory indexes significantly differed as a function of sex and genotype in both NOR and OiP tasks (sex*genotype: OiP F1,64 =12.27, p = 0.0008, NOR F1,64 = 7.11, p = 0.0097; Fig. 1C, D respectively). In the OiP task, dKI females performed at the chance level and significantly worse than WT females and dKI males. WT of both sexes and dKI males performed similarly well and above chance level (p < 0.001). In the NOR task, the memory index of dKI females was also significantly lower than that of WT females and dKI males. At this age, all four groups performed above the chance level (dKI females: p < 0.01; other groups: p < 0.0001).

At the age of 4 months, there was no sex and genotype interaction, but dKI mice were globally impaired compared to WT mice (genotype: OiP F1,63 = 69.65, p < 0.0001, NOR F1,63 = 74.49, p < 0.0001; Fig. 1E, F respectively). A significant genotype difference was also present within each sex for both tasks. WT females and males performed better than their dKI counterparts and above the chance level (p < 0.0001). The memory index of dKI females slightly differed from chance level (p < <0.05), whereas dKI males performed at chance level. To sum up these behavioral results, memory performances in both tasks are affected in dKI females at the age of 2 and 4 months and in dKI males at 4 months compared to their WT counterparts.

### Complex sex and genotype (dKI vs. WT) interactions in Default Mode Network patterns

Group ICA was employed to decompose the rs-fMRI signal into spatially independent components, allowing the identification of distinct functional networks across the brain. We tested multiple component numbers (15, 30, 50, and 100) for ICA. Among these, the 30 and 50 models were selected for further inter-group analysis, as they provided representative mapping of DMN-like patterns (Fig. 2). The 30-component ICA approach revealed a single DMN-like component (Fig. 2A), while the 50-component ICA approach segregated the DMN into three adjacent subnetworks corresponding to its rostral, medial and caudal parts (Fig. 2B). These representative DMN components included canonical regions such as the orbitofrontal cortex, medial prefrontal cortex (mPFC), anterior cingulate cortex (ACA, rostral and medial parts), RSP. Additional recruitment was observed in parts of HIP, somatosensory (SS) and ENT, pallidum (PAL; part of the nucleus basalis), and striatum, as shown in Fig. 2.

**Figure 2.**
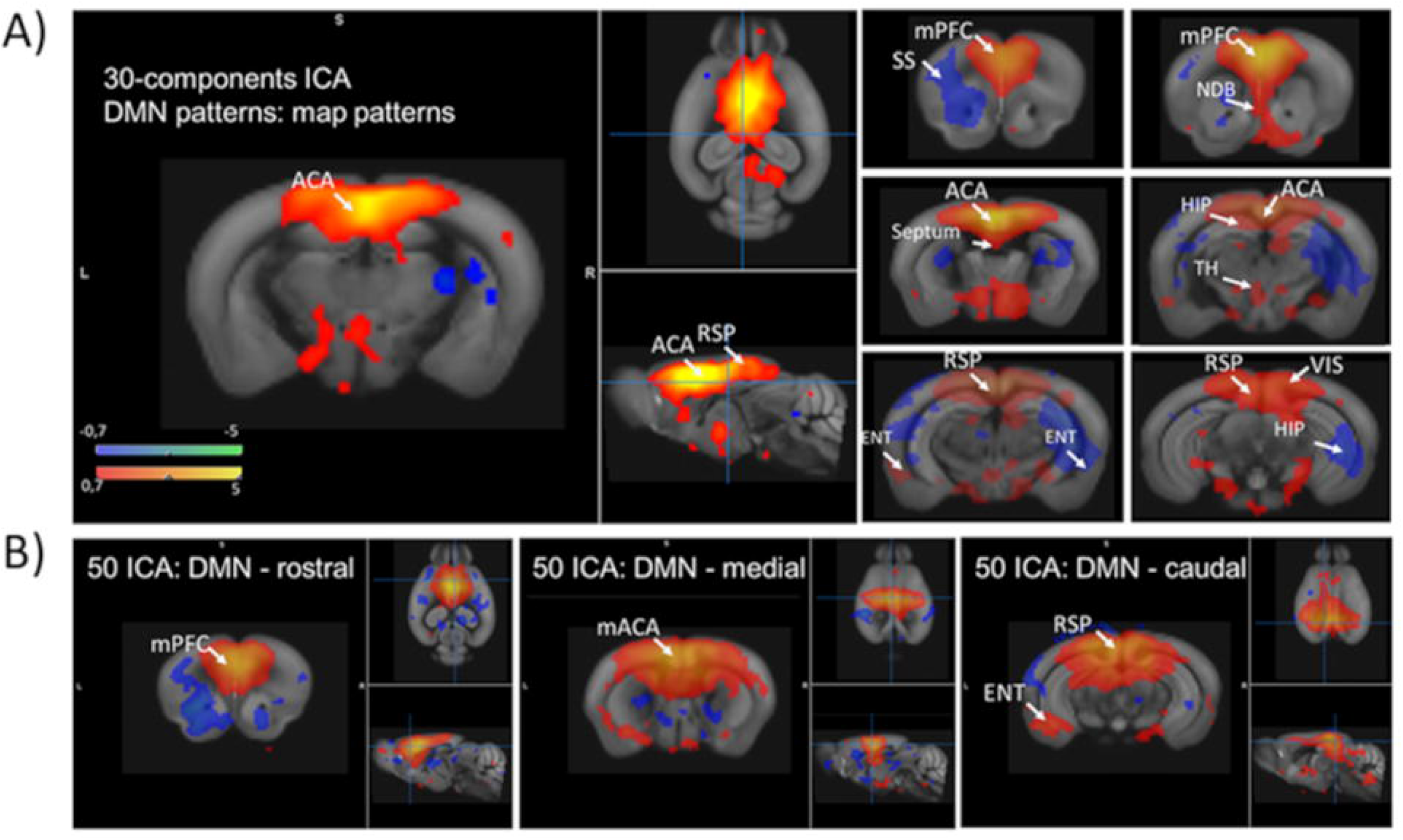
DMN-like patterns. derived with 30-components ICA model (A) and the corresponding segregation in 3 subnetworks using 50-components ICA model. Spatial maps represent the average pattern of this component including positive (red) and negative (blue) correlations calculated voxel-wise with the mean time course of the component. Areas included are: mPFC, ACA, RSP – canonical regions of DMN – but also parts of septum, HIP, TH, and ENT cortex. Abbreviations are listed in Supplemental Table 1.

We selected the 30 ICA-derived DMN components and generated group incidence maps (Supplemental Fig. S2) which demonstrated consistent connectivity across animals. This robustness is evidenced by the incidence maps observed in the dKI groups (Figs. S2A and C) and WT groups (Figs. S2B and D). A full-factorial statistical analysis of this component revealed significant sex- and genotype-specific variations in DMN connectivity. **Sex differences** were evident at 2 months of age in both WT and dKI mice (Fig. 3A and B). Female WT mice exhibited within DMN hypoconnectivity or hyposynchrony compared to male WT mice in regions overlapping the mPFC, medial ACA (mACA), and small clusters within the motor (MO) and SS cortices. This was contrasted by hyperconnectivity in a small cluster within caudal RSP and superior colliculus relative to the WT male counterparts (Fig. 3A). Female dKI mice showed within DMN hyperconnectivity/hypersynchrony compared to male DKI DMN patterns in regions overlapping mACA/MO, SS cortices and dorsal and ventral hippocampal areas. This was contrasted by a small left mPFC area, which exhibited lower FC within DMN compared to male dKI counterparts (Fig. 3B).

**Figure 3.**
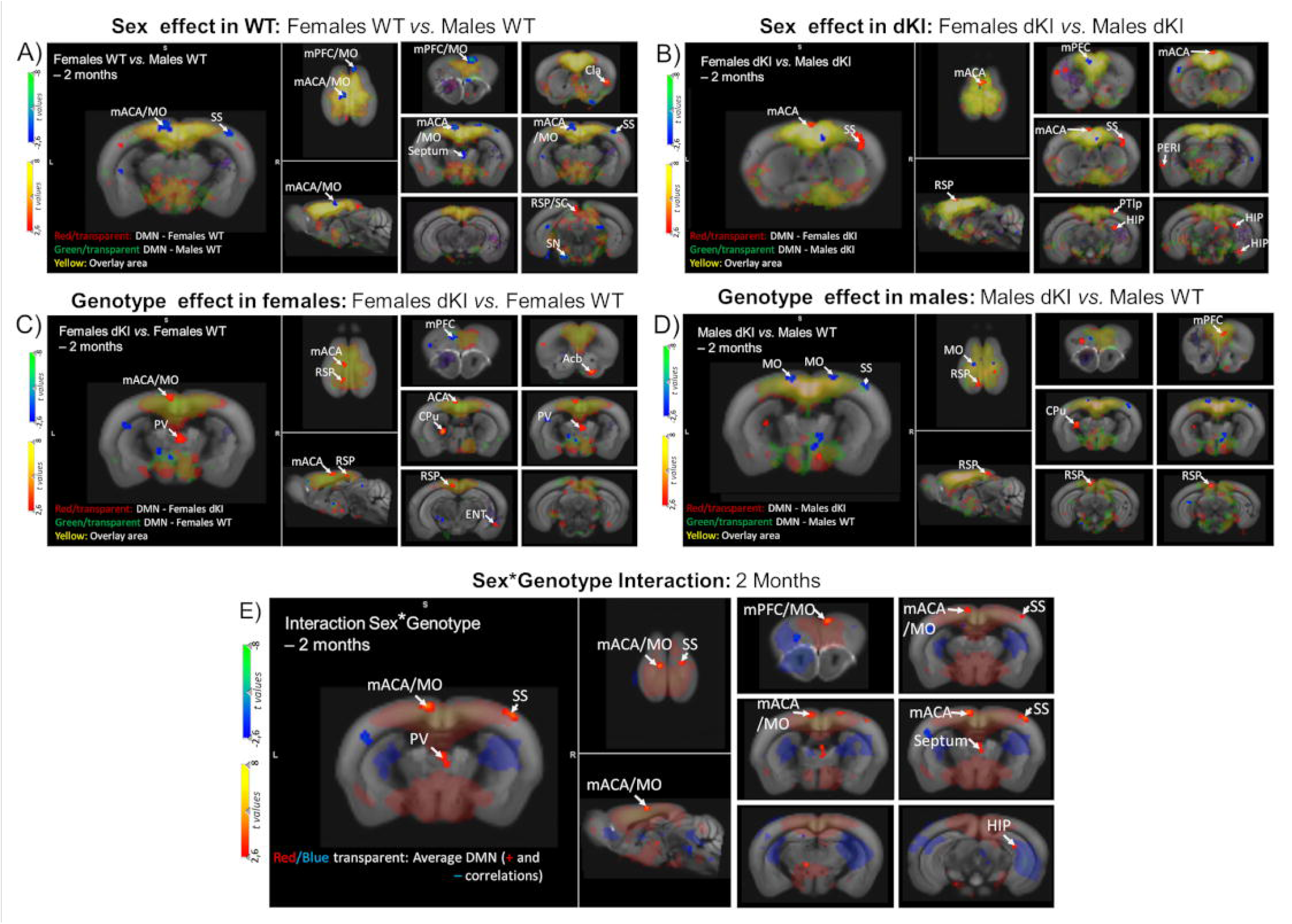
Full-factorial statistical comparison of ICA-derived DMN at 2 months of age. (**A-D**). Subject-specific DMN maps were included in the analysis and inter-group analysis was carried out. Sex effects (females vs. males) are presented separately for WT (A) and dKI mice (B). Genotype effects (dKI vs. WT) are presented for females (C) and males (D). Sex genotype interactions are given in panel F. Statistical results (red, blue - t values, thresholded for p<0.02, uncorr., with a minimal cluster size of 20 voxels) are overlaid on average maps of the DMN components corresponding to the compared groups (yellow – in transparent on the Allen Mouse Brain Atlas Template). A) **Females WT vs. Males WT comparison.** Scale bars: Red/yellow indicate areas of stronger connectivity in the areas (higher synchrony) within DMN in Females WT compared to Males WT. **Blue/green** indicate areas of lower connectivity within DMN in Females WT compared to Males WT. B) **Females dKI vs. Males dKI comparison.** Scale bars: Red/yellow indicate areas of stronger connectivity in the areas (higher synchrony) within DMN in Females dKI compared to Males dKI. **Blue/green** indicate areas of lower connectivity within DMN in Females dKI compared to Males dKI. C) **Females dKI vs. Females WT comparison.** Scale bars: Red/yellow indicate areas of stronger connectivity in the areas (higher synchrony) within DMN in Females dKI compared to Females WT. **Blue/green** indicate areas of lower connectivity within DMN in Females dKI compared to Females WT. D) **Males dKI vs. Males WT comparison.** Scale bars: Red/yellow indicate areas of stronger connectivity in the areas (higher synchrony) within DMN in Males dKI compared to Males WT. **Blue/green** indicate areas of lower connectivity within DMN in Males dKI compared to MalesWT. E) **Sex-by-Genotype interaction:** Red/yellow indicates more pronounced genotype-related differences in females. **Blue/green** indicate areas with more pronounced genotype-related differences in males. Experimental groups are the following: WT females (n=17), dKI females (n=17), WT males (n=16), dKI males (n=17). Abbreviations are listed in Supplemental Table 1.

**Genotype effects** were assessed by comparing dKI and WT groups within each sex (Fig. 3C, D). Female dKI mice showed within DMN hypoconnectivity in a small cluster of the left mPFC but demonstrated hyperconnectivity in regions spanning parts of the medial ACA/MO areas, clusters within RSP, the right ENT, and caudate putamen and paraventricular (PV) thalamic regions compared to WT females (Fig. 3C). Conversely, male dKI mice displayed a small cluster of mPFC increased connectivity within DMN and minimal or no difference in mACA compared to WT males (Fig. 3D). Genotype effects were also notable in the ENT, where female dKI mice exhibited increased DMN connectivity relative to WT females. However, no significant differences in ENT connectivity within DMN could be observed between male dKI and male WT mice (Fig. 3C and D). Both female and male dKI mice showed increased connectivity of RSP areas within the DMN when compared to their respective WT controls (Fig. 3C and D). **Sex-by-genotype interactions** at 2 months of age (Fig. 3D) revealed that regions within DMN overlapping the ACA/MO, SS, PV, and HIP exhibited more pronounced genotype differences in females, driven by hyperconnectivity in dKI females within these DMN overlapping regions.

Overall, these findings demonstrate that genotype and sex significantly influence FC within the DMN and its interactions with memory-associated regions in the dKI mouse model at 2 months of age. By exploring the DMN’s organization our analysis reveals the ACA and RSP as critical nodes for these sex and/or genotype modulations. These sex-by-genotype interactions were markedly reduced at 4 months and remained relevant only for RSP (Supplemental Fig. S3). Given that the RSP serves as a pivotal hub within the DMN and is strongly implicated in memory consolidation and spatial navigation, we selected this region in a seed-based approach to explore its positioning within the broader brain network in dKI mice, focusing on analysis of data from mice of 2 months of age.

### Sex and Genotype-specific interaction between DMN and memory processing key nodes

Within the functional network of the RSP, hyperconnectivity with the **ENT** was identified as a prominent feature (Fig. 4A). This hyperconnectivity was observed across multiple comparisons: WT females relative to WT males (Fig. 4A), dKI females relative to dKI males (Fig. 4B), and dKI females relative to WT females (Fig. 4C). However, no significant differences were detected in dKI males compared to WT males (Fig. 4D). These results highlight a **strong sex effect on the RSP-ENT connectivity axis.**

**Figure 4.**
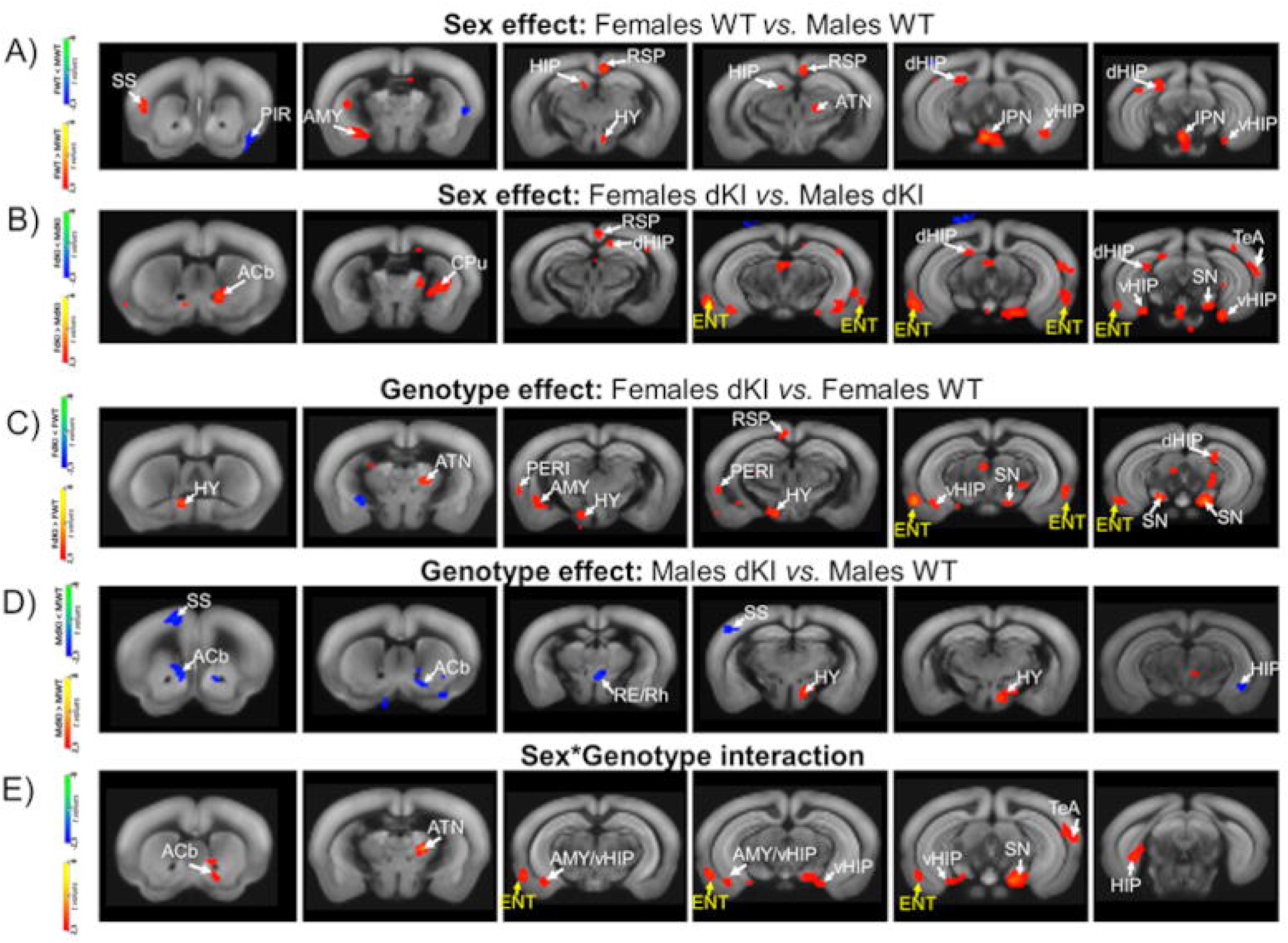
Inter-group statistical differences in RSP functional connectivity (RSP FC) in 2-month-old mice. (A) Statistical maps show **sex-specific differences in RSP FC in wild-type (WT) mice** (female WT vs. male WT). Red/yellow areas show stronger RSP FC in females WT compared to males WT, while blue/green areas exhibit weaker RSP FC in females WT relative to males WT. (B) Statistical maps show **sex-related differences in the RSP FC in dKI mice** (females dKI *vs.* Males dKI). Red/yellow areas show stronger RSP FC in females dKI compared to males DKI, while blue/green areas exhibit weaker RSP FC in females dKI relative to males dKI. (C) Statistical maps show **genotype-related differences in the RSP FC in female mice** (females dKI *vs.* females WT). Red/yellow areas show stronger RSP FC in females dKI compared to females WT, while blue/green areas exhibit weaker RSP FC in females dKI relative to females WT. (D) Statistical maps show g**enotype-related differences in the RSP FC in male mice** (males dKI *vs.* males WT). Red/yellow areas show stronger RSP FC in males dKI compared to males WT, while blue/green areas exhibit weaker RSP FC in males dKI relative to males WT. F) **Contrast maps showing areas exhibiting significant Genotype*Sex interactions.** Red/yellow indicates more pronounced genotype-related differences in females. **Blue/green** indicate areas with more pronounced genotype-related differences in males. Abbreviations are listed in Table 1. **For all comparisons (A, B, C, D, E) statistical maps were generated** using a full factorial design (sex, genotype, and age as variables) with contrast selection in SPM. **Scale bars indicate** t values displayed for p < 0.02, with a minimum cluster size of 20 voxels. Individual RSP’s functional networks introduced in the statistical comparisons were mapped using an RSP mask as a seed, and Spearman correlation was carried on voxel-wise across the whole brain. Abbreviations are listed in Supplemental Table 1.

At 2 months of age, dKI females also exhibited hyperconnectivity between the RSP and anterior thalamic nucleus (ATN) as well as the substantia nigra (SN) when compared to WT females (Fig. 4C). These findings suggest **genotype-driven alterations** in thalamocortical and midbrain-cortical communication specific to females.

Conversely, male dKI mice displayed distinct hypoconnectivity patterns at 2 months of age, particularly between the RSP and the **reuniens nucleus (RE),** SS, and **HIP** (Fig. 4D). These FC changes indicate that the dKI genotype exerts sex-dependent effects on the RSP functional network.

Interestingly, **RSP - HIP** connectivity revealed **contrasting sex and genotype effects**. Female WT mice exhibited stronger connectivity between RSP and areas of the dorsal HIP (dHIP) and ventral HIP (vHIP) compared to males (Fig. 4A) This higher connectivity was further enhanced in dKI females compared to WT females (Fig. 4C) for both RSP to dHIP and RSP to vHIP. In contrast, dKI males displayed decreased RSP-vHIP connectivity, highlighting genotype effects that are sex-specific in the RSP-HIP connectivity.

**Sex-by-genotype interactions** (Fig. 4E) revealed that RSP connectivity with multiple subcortical and cortical regions, including the accumbens (ACb), ATN, vHIP, SN, and particularly ENT cortex exhibited more pronounced genotype-related differences in females. These effects were primarily driven by the hyperconnectivity of RSP with these regions in females dKI.

In summary, these findings underscore a pronounced sex-driven pattern of hyperconnectivity between the RSP and memory-associated regions, such as the ENT and HIP, in dKI female mice. The observed hyperconnectivity suggests a potential early mechanism underlying disrupted communication between DMN and memory-related networks in this model of early-onset AD. In contrast, decreased patterns of FC observed in male dKI mice (e.g., RSP-RE, RSP-SS, RSP-HIP) indicate sex-divergent connectivity changes. These findings highlight distinct DMN–memory hub interactions that fundamentally differ between males and females at 2 months of age.

## Discussion

In this study, we used the *App^NL-F^/MAPT* dKI mouse model (Saito *et al*., 2019) to investigate sex-dependent connectivity patterns of DMN during the preclinical phase of AD. Behavioral tests at 2 and 4 months of age, evaluating short-term associative and long-term recognition memory, were combined with rs-fMRI to map cerebral network connectivity. Our findings revealed that memory deficits emerged earlier in dKI females compared to males, a phenomenon often observed in human AD studies (Lin *et al*., 2015; Koran *et al*., 2017). This earlier onset of deficits in females strengthens the validity of the dKI mouse model for studying the sex-dependent emergence of AD features. Deficits occurring in both tasks as early as 2 months of age in dKI females were accompanied by distinct FC changes within the DMN and between its hubs and memory-processing nodes distributed across the brain. Female dKI mice displayed hypersynchronous resting-state activity within the DMN, particularly in the RSP, a critical DMN hub in mice (Karatas *et al*., 2021; Mandino *et al*., 2021; Whitesell *et al*., 2021). This RSP hypersynchrony extended beyond canonical DMN regions, engaging key memory-processing areas such as the perirhinal (PERI) and ENT cortices, HIP, amygdala, thalamic nuclei (PV and ATN), and dopaminergic areas such as SN. These patterns emerged as significant sex*genotype connectivity interactions and point towards circuitry dysfunctions as early markers of sex-dependent cognitive deficits. A small cluster of frontal DMN part (left mPFC) exhibited, however, hyposynchronous activity within DMN, indicating a complex, region-specific, and potentially lateralized modulation of connectivity within this network in memory-deficient dKI females. Meanwhile, at 2 months of age, male dKI mice performed comparably to their WT counterparts, while memory deficits were found at 4 months. Notably, despite the absence of overt memory impairments at 2 months, subtle FC remodelling was already evident in male dKI mice compared to WT males. Within the DMN ICA component, we found hypersynchronous rs-fMRI signal in a limited area of the caudal RSP and a small area of the right prefrontal cortex (PFC). However, seed analysis revealed that RSP connectivity across the whole brain diverged from hypersynchronous patterns observed in female dKI. Notably, RSP showed reduced FC with HIP, RE/rhomboid thalamic nuclei, but also somato-motor cortical areas and ACb nuclei. These findings align with the c-Fos-based ex vivo imaging study by Borcuk et al. (2022), which reported disrupted RSP connectivity with the HIP and medial temporal lobe cortices (ENT and PERI) in 4-month-old male dKI mice (Borcuk *et al.,* 2022). Our seed-based FC analysis that similarly revealed already at 2 months RSP–HIP hypoconnectivity in male dKI mice compared to WT males, highlights a pattern of network dysfunction that precedes the onset of cognitive deficits. Such network remodelling, emerging prior any cognitive deficits aligns with our previous observations in Ty22-Tau AD male mice (Degiorgis *et al*., 2020) and is consistent with results from other studies showing early FC modifications using different models of AD (Shah *et al*., 2016a; Latif-Hernandez *et al*., 2019). These findings support the hypothesis that connectivity alterations, particularly within the DMN, may serve as predictive hallmarks of AD clinical manifestation and memory performance (Ju *et al*., 2023). Our results further demonstrate that this remodelling exhibits a complex, sex-specific pattern.

### DMN hypersynchrony as a marker of preclinical AD and female network vulnerability

Our rs-fMRI data from females *App^NL-F^/MAPT* dKI mice share strong similarities with the findings of Shah et al. in female *App^NL-F^* knock-in (KI) mice (Shah *et al*., 2018, 2022), where hypersynchrony of DMN hubs and particularly cingulate cortex hypersynchrony emerged as cardinal features of early pathology. In 3-month-old female *App^NL-F^* KI mice, spatial reversal learning defects were associated with hypersynchronous FC of the HIP, ACA, caudate-putamen, and, more largely the DMN (Shah *et al*., 2018). In a subsequent translational imaging study in both mice and humans, Shah et al. (2022) demonstrated that DMN alterations, characterized by hyperconnectivity in the cingulate cortex, occur even earlier, between 1.5 and 3 months of age in female *App^NL-F^* KI mice (Shah *et al*., 2022). This disruption was accompanied by neuronal hyperactivity and reduced astrocyte calcium signalling. Importantly, local restoration of regulatory activity of astrocytes reversed neuronal hyperactivity, cingulate hyperconnectivity, and behavior deficits, supporting a model where cingulate astrocytes play a pivotal role in mediating initial features of AD and clinically relevant phenotypes. The authors synergistically showed that increased FC in the cingulate cortex is not only a feature of cognitively normal individuals at risk for AD (Kucikova *et al*., 2021) but also predicts amyloid-positron emission tomography (PET) positivity years before clinical symptoms, positioning this increased FC as one of the earliest biomarkers of incipient AD (Jack *et al*., 2017; Shah *et al*., 2022). Regarding amyloid pathology, *App^NL-F^*KI mice exhibit an increased Aβ42/40 ratio by 3 months of age and initial amyloid plaque deposition around 6 months (Saito et al., 2014). Similarly, at 2 and 4 months, *App^NL-F^/MAPT* dKI mice also demonstrate an increased Aβ42/40 ratio (Borcuk *et al.,* 2022), consistent with observations in *App^NL-F^* KI mice, and no amyloid plaque deposition until late progressive accumulation throughout their lifespan (Saito *et al*., 2019; Islam *et al*., 2023; Aguilera *et al*., 2024). Additionally, in the *App^NL−G−F^* KI model with more severe pathology, Latif-Hernandez et al. (2019) observed subtle behavioral changes and increased prefrontal-hippocampal network synchronicity in females before Aβ plaque deposition (Latif-Hernandez *et al*., 2019). Using rs-fMRI, they reported hypersynchronized activity between DMN and memory-related regions, particularly the PFC-HIP network, as early as 3 months of age. In an associated study on *App^NL-G-F^* mice at 3–4 months of age, synaptic impairment, as indicated by a deficit in long-term potentiation was demonstrated in the PFC of these mice (Latif-Hernandez *et al*., 2020). This impairment became more pronounced and extended to the hippocampus by 6–8 months. In a longitudinal study in the same *App^NL-G-F^* model, Morrissey et al. (2023) also showed higher interhemispheric connectivity between hippocampal subregions, followed by a later decrease, in *App^NL-G-F^* female mice compared to WT females, as pathology progresses (Morrissey *et al*., 2022). Our data aligns with the observed hypersynchrony between DMN nodes and HIP; however, the main axes of hyperconnectivity in our female *App^NL-F^/MAPT* dKI are RSP-ENT and RSP-HIP. This pattern was also evident as a sex effect in WT mice, with females WT showing stronger connectivity than males WT at 2 months. However, the hyperconnectivity effect along RSP-vHIP / RSP-ENT is exacerbated in our female *App^NL-F^/MAPT dKI* mice and emerges as a strong sex*genotype interaction at 2 months. *This marked hyperconnectivity between key DMN hubs and memory nodes supports the idea of its relevance as an early predictor and marker of AD initiation. It stands as well as a potential discriminative signature of early vulnerability in the female population*.

### Sex Differences in DMN Modulation and Memory Processing

In the context of human research, the sex dimorphism of DMN FC and its interaction with memory processing nodes during aging, which could clarify women’s higher vulnerability to AD, is poorly understood. Limited studies suggest that aging women exhibit increased DMN connectivity compared with men, predominantly in posterior nodes (Biswal *et al*., 2010; Scheinost *et al*., 2015; Ferretti *et al*., 2018; Ritchie *et al*., 2018) and mostly during perimenopausal decades (Ficek-Tani *et al*., 2023a). Additionally, it was shown that women rely more on the DMN than men for short-term memory performance (Natu *et al*., 2019; Kang *et al*., 2021; Vanneste *et al*., 2021; Ficek-Tani *et al*., 2023a). Ju and colleagues (2023) applied connectome-based predictive modelling (CPM), a robust linear machine-learning approach to the Lifespan Human Connectome Project-Aging (HCP-A) demonstrated that women and men recruit different circuitry when performing memory tasks, with women relying more on intra-DMN activity and men relying more on visual circuitry (Ju *et al*., 2023). Our comparison of female WT and male WT groups highlights similar patterns of higher dorso-caudal RSP connectivity in females and increased RSP-HIP; suggesting sex as a modulator of DMN connectivity and its interaction with major memory processing hubs. As proposed by Ju et al., the female sex relatively increased reliance on DMN connectivity could lead to “overuse” and vulnerability of this network to AD, as well as earlier and more severe pathologic burden over time.

Furthermore, the hyperconnectivity observed in our dKI females mirrors also findings in human studies, where preclinical AD is associated with increased DMN connectivity in posterior regions (e.g., RSP) and memory hubs (e.g., HIP) (Mormino *et al*., 2011; Ficek-Tani *et al*., 2023b). These patterns are thought to represent either a precursor to amyloid deposition itself (Jones *et al*., 2016; Shah *et al*., 2016b) or, alternatively, a compensatory response to pathology (Qi *et al*., 2010). Furthermore, within DMN, RSP and posterior cingulate cortex (PCC) were found to be characterized by differential FC patterns. These patterns are characterized by hub-specific interactions with memory and attention scores in prodromal AD, compared to cognitively normal individuals, possibly reflecting compensatory mechanisms for RSC and neurodegenerative processes for PCC (Dillen *et al*., 2016, 2017). Jack et al. (2013) provided evidence for a preclinical phase of AD in Aβ-negative individuals that had increased posterior DMN connectivity before becoming Aβ-positive at follow-up (Jack *et al*., 2013, 2017). Similarly, increased FC has been observed in individuals carrying the APOEε4 allele, a known risk factor for AD (Filippini *et al*., 2009; Sheline *et al*., 2010). Of interest, the female sex is associated with a greater pathology burden in the early onset AD compared to the male sex (Nemes *et al*., 2023). Moreover, women with APOEε4 allele and amyloid PET positivity showed higher connectivity between anterior and posterior DMN, than their ε4-counterparts, while no differences were observed in men.

Bero and colleagues demonstrated in APP transgenic mouse models that patterns of bilateral RSP hyper-connectivity at a young age correlated with amyloid plaque distribution later in life (Bero *et al*., 2012). Other studies combining rs-fMRI and PET imaging in humans (Schultz *et al*., 2017a) have shown that elevated Aβ alone is linked to DMN hyperconnectivity, while the presence of both Aβ and Tau pathologies is associated with DMN hypoconnectivity. Similarly, young AD transgenic mice at an early soluble Aβ stage showed hypersynchronisation across DMN regions, while overall hypoconnectivity coincided with a widespread deposition of Aβ plaques at an older age (Ben-Nejma *et al*., 2019). Tau pathology accumulates early in the medial temporal lobe areas - including ENT and PERI cortices, regions strongly connected to cortical DMN (Ward *et al*., 2014). Relying on the interplay of DMN and ENT/PERI activity, episodic memory deficits are typically the most salient cognitive indicators of early AD (Tromp *et al*., 2015). Interestingly, during the MCI phase in humans, sex differences in FC have also been reported. Males show higher global connectivity, while females exhibit greater within-network integration (Blujus, 2021).

The *App^NL-F^/MAPT* dKI mice, which incorporate both amyloid and tau pathology at older age, would be particularly valuable for studying the combined effects of these two hallmark proteins of AD and following FC pattern changes from preclinical to more advanced stages.

The sex-divergent patterns that we observe along the RSP-HIP and RSP-ENT connectivity in the *App^NL-F^/MAPT* dKI mice warrant discussion in relationship with the recruitment of these specific pathways in memory tasks we used. However, as both tasks were impaired together in deficient male and female groups, it precluded a specific association between patterns of FC alteration and deficits in one of these tasks. Nevertheless, the vast majority of regions involved in dKI DMN abnormal connectivity (e.g., mPFC, RSP, HIP, ENT, PERI, RE/RH, Septum, Hy) were shown to be involved in OiP associative memory and/or in long-term NOR in studies using invasive activation or inactivation approaches in rodents (for review see Mathis and Lecourtier, 2017; Chao *et al*., 2022). Interactions of the RSP with PERI, ENT, mPFC, and even the ATN are required for long-term NOR (de Landeta *et al*., 2020, 2021). RSC appears as a key region or OiP associative memory, although the role of its connections with other regions has been poorly investigated (Vann and Aggleton, 2002). The mPFC-HIP pathway is crucially involved in short-term OiP memory and in NOR consolidation (Barker and Warburton, 2015; Tuscher *et al*., 2018).

### Longitudinal DMN remodelling in AD Progression

One of the most compelling observations from both animal and human studies literature is that DMN alterations in early AD are not static (Zerbi *et al*., 2014; Schultz *et al*., 2017a; Ben-Nejma *et al*., 2019; Morrissey *et al*., 2022; Mandino *et al*., 2024). Instead, a transition from a state of hyperconnectivity to hypoconnectivity is frequently described as the disease advances (Brier *et al*., 2012; Schultz *et al*., 2017b). Staffaroni and colleagues recently showed that DMN FC in healthy aging follows a trajectory of increases in connectivity early in the aging process, with more rapid declines in older age. Connectivity patterns within DMN were found as a marker of episodic memory performance, even among cognitively healthy older subjects (Staffaroni *et al*., 2018). This trajectory, however, can be nuanced by many factors, including sex, genetics, and strongly by amyloid and tau accumulations, as the specific impact of Aβ and tau on DMN connectivity depends on the stage of pathology (Buckner *et al*., 2005). In a longitudinal study, Mandino and colleagues investigated the evolution of connectivity changes at 4, 6, and 9 months of age in *App^NL-G-F^/MAPT* dKI mice, comparatively to healthy aging WT mice, in a mixed-sex cohort. They found a widespread increase in FC in the WT group with aging, which persisted until the study endpoint. In contrast, while 4-month-old AD mice were similar to WT controls, significant DMN hypoconnectivity emerged in the AD mice by 6 months and became more pronounced by 9 months. Although direct comparisons with our younger cohort of *App^NL-F^/MAPT* dKI mice are challenging, their findings at 4 months resemble more to the results we observed in the male dKI cohort.

For further translation of imaging biomarkers, it is therefore important to acknowledge the variability in findings across different animal models. Some models show early hypersynchronization in the hippocampus (Morrissey *et al*., 2022), while others show more widespread changes (van den Berg *et al*., 2022). These differences can stem from the type of model (transgenic vs. knock-in), specific mutations used, the timing of transgene expression, and last but not least - the sex on which the research was conducted – all of which should be interpreted with caution. Methodological variations, such as c-Fos, EEG, or rs-fMRI for data collection, and differences in analytical algorithms further contribute to variability but also bring complementary knowledge on the mechanisms of brain communication and response in early AD. For instance, static FC measures from our study, revealing increased connectivity within “typical” functional network patterns such as DMN, may also reflect an over-representation of specific brain states resulting from a loss of FC dynamics. This highlights the importance of further investigating the dynamic patterns of FC in both male and female *App^NL-F^/MAPT* dKI mice. Understanding the spatiotemporal dynamics of brain patterns in our model would improve the characterization of network disruptions in preclinical AD and further refine the sex-specific signatures. Such approaches were recently applied to reveal the link between static and dynamic brain functional network connectivity and genetic risk of AD (Sendi *et al*., 2023) and showed that genetic risk synergistically affects the whole-brain static and dynamic FC in women but not men.

In conclusion, our data in *App^NL-F^/MAPT* dKI mice point towards an increased dorso-caudal hypersynchrony of the RSP within the DMN during the preclinical stages of AD, observed in both sexes but with distinct sex-specific interactions with memory hubs. A female-specific pattern of hypersynchrony between RSP and the HIP/ENT was evidenced at 2 months, coinciding with the onset of cognitive deficits in females. In contrast, male dKI mice show a pattern of hyposynchrony along these connection axes at both 2 and 4 months, particularly along RSP-HIP connectivity. This might reflect a female-specific circuitry vulnerability early in AD or circuitry resilience and compensatory remodeling in males. The hypoconnectivity observed in dKI males raises intriguing questions about its potential protective role in limiting the spread of pathological proteins like tau and amyloid (Wang *et al*., 2024). The presence of tau pathology in the ENT, but not yet in the HIP, in 4-month-old dKI males (Borcuk *et al*., 2022) supports the idea that ENT plays a pivotal role in early AD pathology. Structural differences between sexes may further contribute to observed functional patterns, emphasizing the importance of integrative studies that combine structural, functional, histological, and molecular data. Our present results underscore the multifactorial influences shaping DMN connectivity and its interactions with memory processing hubs in AD. They also emphasize the critical importance of incorporating sex-specific analyses in preclinical AD research to unravel the nuanced mechanisms underlying disease progression.

## Acknowledgments

This work was supported by funding from the Fondation pour la Recherche Médicale (FRM – ALZ20192009643), the Université de Strasbourg, and the Centre National de la Recherche Scientifique (CNRS). We extend our warmest thanks to Drs. Shoko Hashimoto, Takaomi C. Saido, and Takashi Saito, as well as the RIKEN BioResource Center (Japan), for providing the **App^NL-F^** and **MAPT** founders for establishing our dKI colony. We are also grateful to the Strasbourg High-Performance Computing (HPC) Center for providing computational resources with MRI analyses, the MRI IRIS imaging facility (Imaging, Robotics, and Innovation in Health) of the ICube Lab (UMR 7357) for access to the MRI system facilities. We thank Marie-Dominique Marinnutti and Céline Héraud for their invaluable support in experimental organization, and Olivier Bildstein for his dedicated care of our mice.

