## Supplemental Fig. S1 for "Exploring sex-specific alterations in early Alzheimer’s disease using network MRI analyses"

### Slide 1
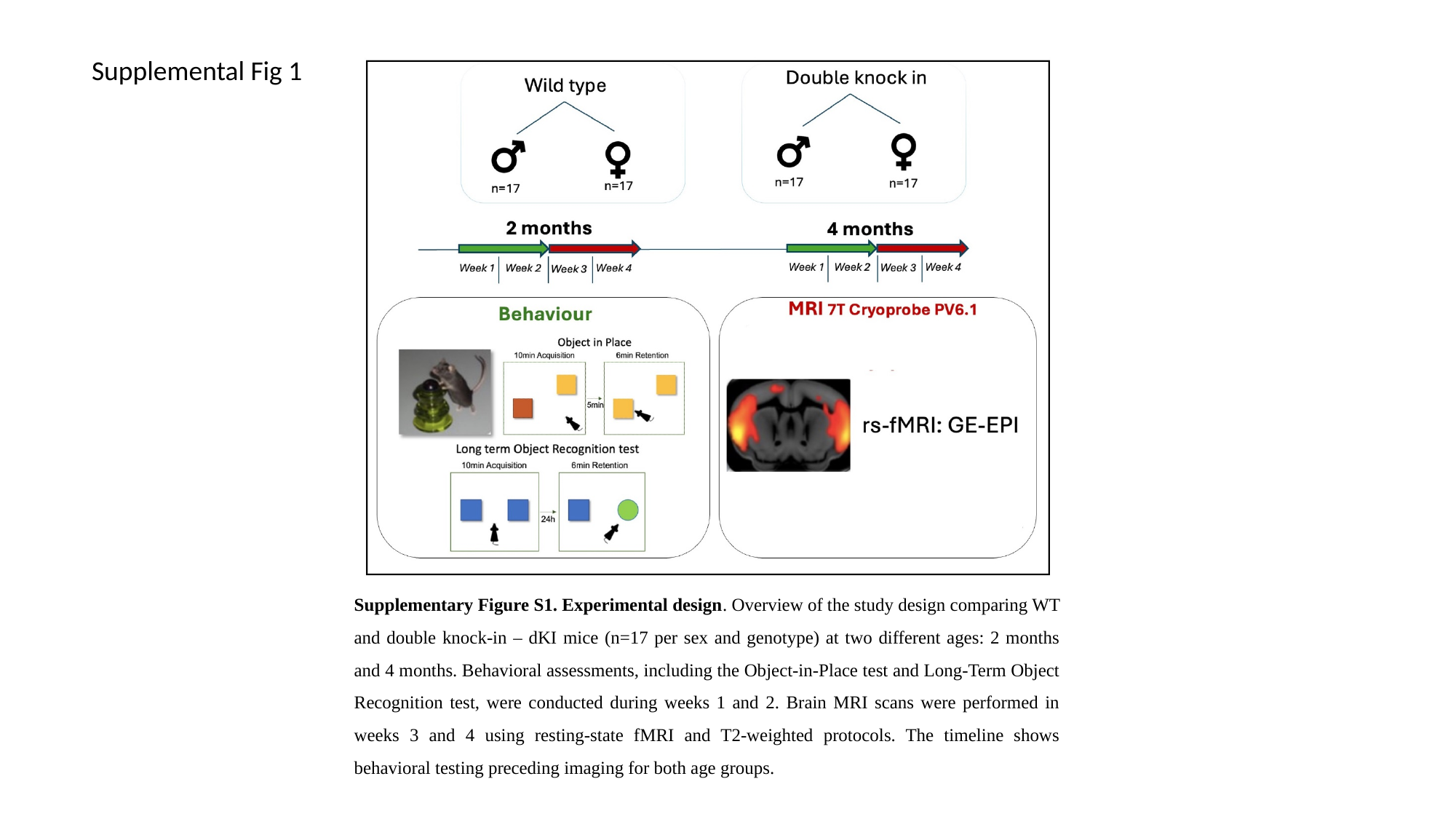

Supplemental Fig 1
Supplementary Figure S1. Experimental design. Overview of the study design comparing WT and double knock-in – dKI mice (n=17 per sex and genotype) at two different ages: 2 months and 4 months. Behavioral assessments, including the Object-in-Place test and Long-Term Object Recognition test, were conducted during weeks 1 and 2. Brain MRI scans were performed in weeks 3 and 4 using resting-state fMRI and T2-weighted protocols. The timeline shows behavioral testing preceding imaging for both age groups.
