## Supplemental Fig. S2 for "Exploring sex-specific alterations in early Alzheimer’s disease using network MRI analyses"

### Slide 1
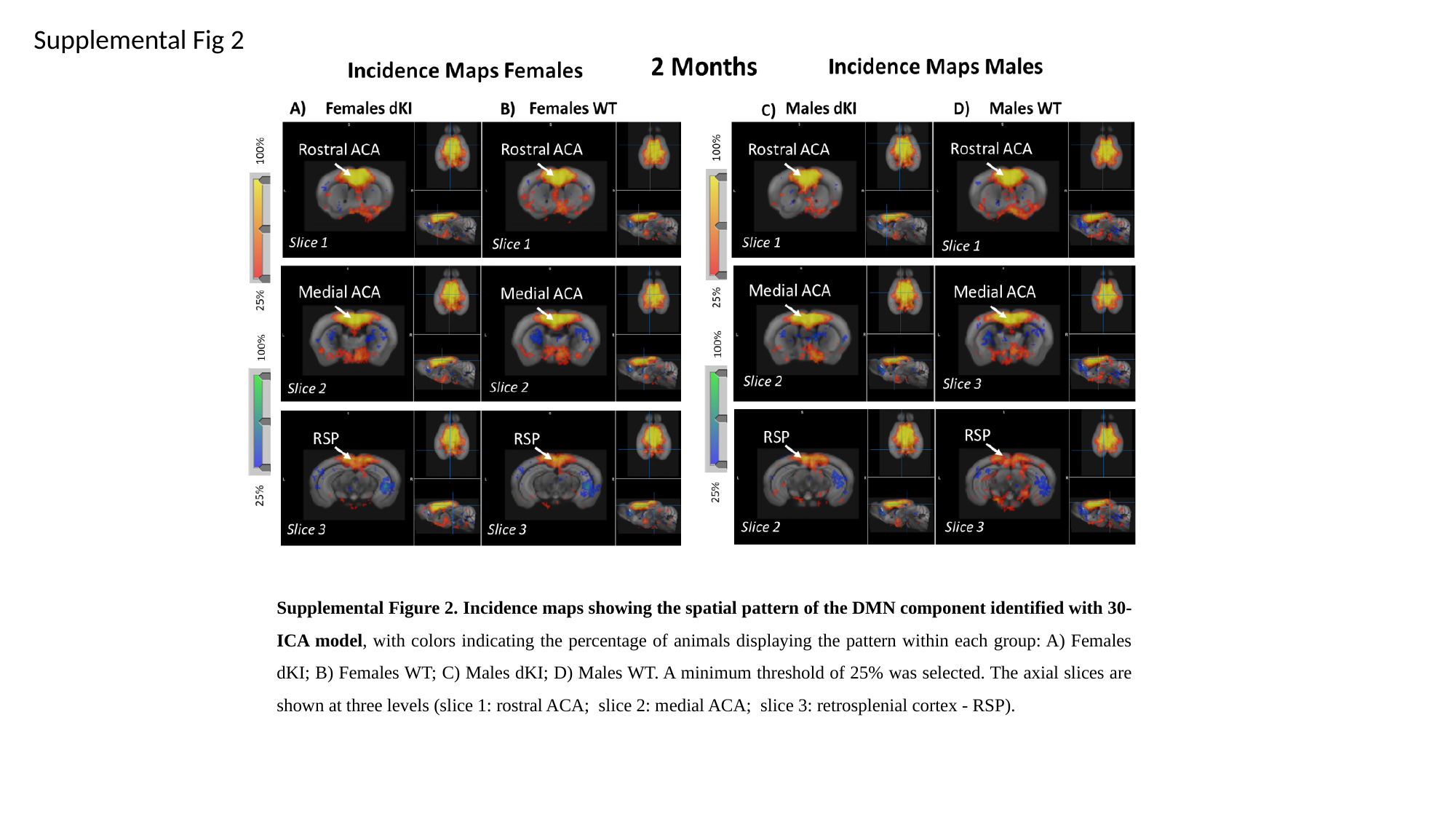

Supplemental Fig 2
Supplemental Figure 2. Incidence maps showing the spatial pattern of the DMN component identified with 30-ICA model, with colors indicating the percentage of animals displaying the pattern within each group: A) Females dKI; B) Females WT; C) Males dKI; D) Males WT. A minimum threshold of 25% was selected. The axial slices are shown at three levels (slice 1: rostral ACA; slice 2: medial ACA; slice 3: retrosplenial cortex - RSP).
