## Supplemental Fig. S3 for "Exploring sex-specific alterations in early Alzheimer’s disease using network MRI analyses"

### Slide 1
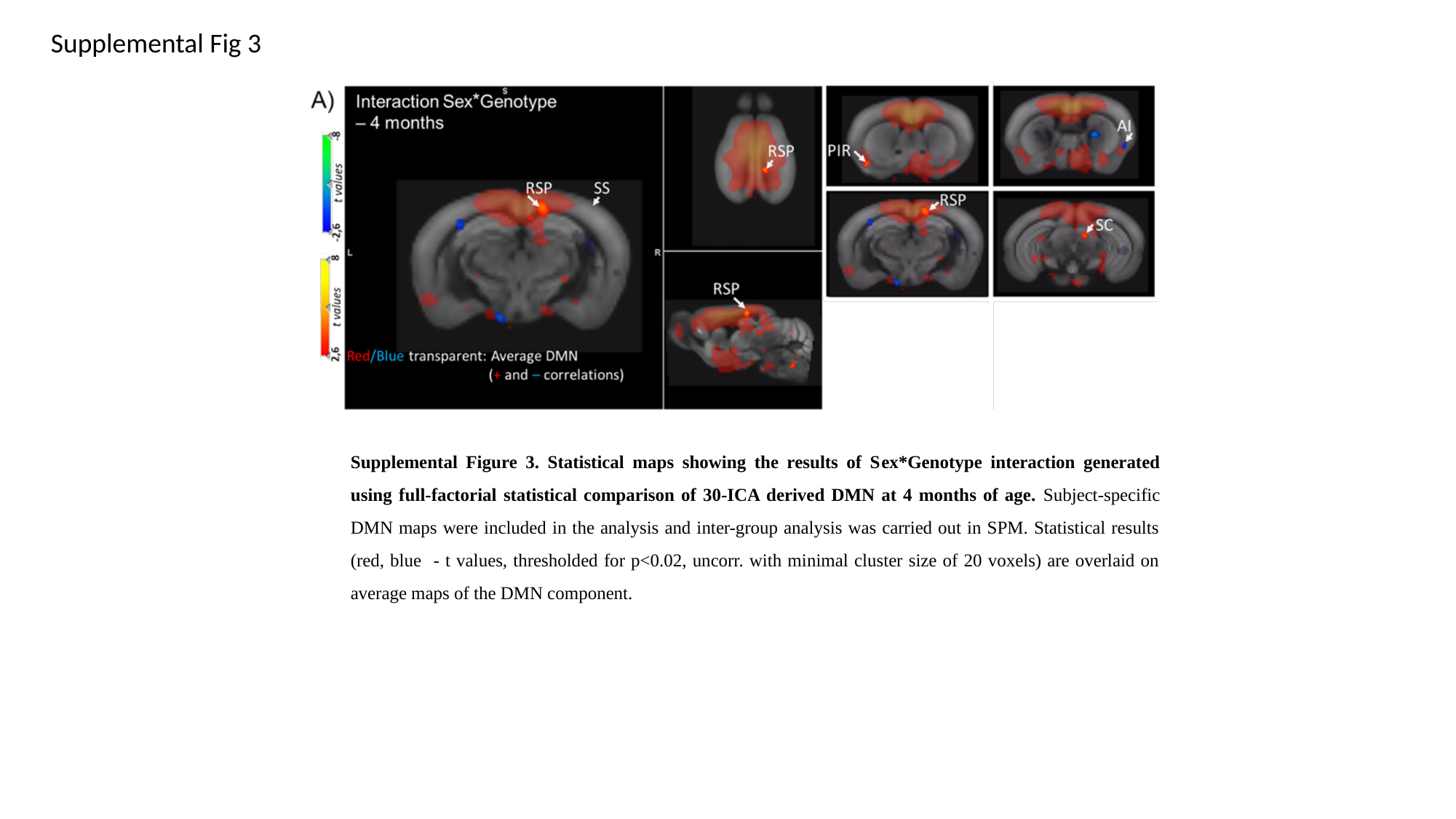

Supplemental Fig 3
Supplemental Figure 3. Statistical maps showing the results of Sex*Genotype interaction generated using full-factorial statistical comparison of 30-ICA derived DMN at 4 months of age. Subject-specific DMN maps were included in the analysis and inter-group analysis was carried out in SPM. Statistical results (red, blue - t values, thresholded for p<0.02, uncorr. with minimal cluster size of 20 voxels) are overlaid on average maps of the DMN component.
