## Supplementary figures and images for "Exploring sex-specific alterations in early Alzheimer’s disease using network MRI analyses"

### Supplemental Table S1

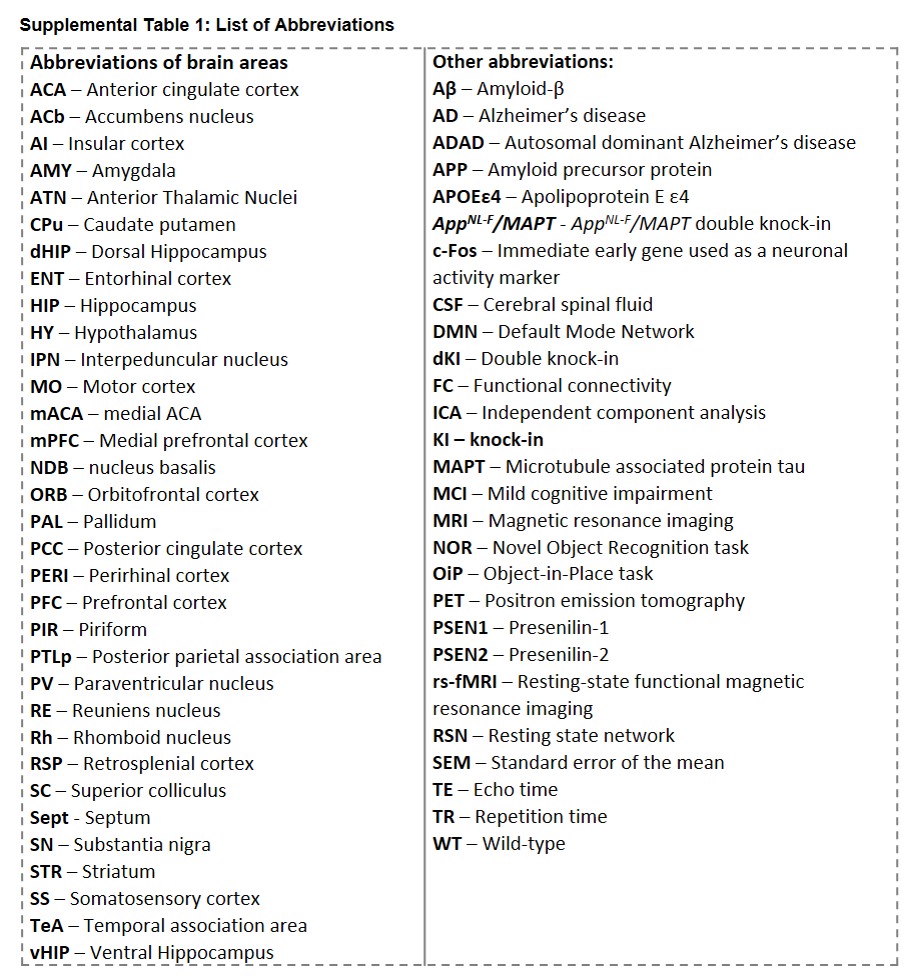
